# Neutralization of the anthrax toxin by antibody-mediated stapling of its membrane penetrating loop

**DOI:** 10.1101/2021.04.18.440036

**Authors:** Fabian Hoelzgen, Ran Zalk, Ron Alcalay, Sagit Cohen Schwartz, Gianpiero Garau, Anat Shahar, Ohad Mazor, Gabriel A. Frank

## Abstract

Anthrax infection is associated with severe illness and high mortality. Protective antigen (PA) is the central component of the anthrax toxin, which is the main virulent factor of anthrax. Upon endocytosis, PA opens a pore in the membranes of endosomes, through which the toxin’s cytotoxic enzymes are extruded. The PA pore is formed by a cooperative conformational change where PA’s membrane-penetrating loops associate, forming a hydrophobic rim that pierces the membrane. Due to its crucial role in anthrax progression, PA is an important target of monoclonal antibodies-based therapy. cAb29 is a highly effective neutralizing antibody against PA. We determined the cryo-EM structure of PA in complex with the Fab portion of cAb29. We found that cAb29 neutralizes the toxin by clamping the membrane-penetrating loop of PA to a static region on PA’s surface, thereby preventing pore formation. Therefore, our results provide the structural basis for the antibody-based neutralization of PA and bring to focus the membrane-penetrating loop of PA as a target for the development of better anti-anthrax vaccines.

## Introduction

Anthrax is a life-threatening infectious disease caused by the bacteria *Bacillus anthracis*. In some modes of infection, the fatality rate from anthrax can be as high as 80%, even with antibiotic treatment [1]. The release of large frozen deposits of the *B. anthracis*, due to thawing of permafrost ground [2], and its relative ease of use as a bioterror agent [3] resurfaced the need for an effective anthrax treatment.

The main virulent factor of *B. Anthracis* is its toxin. The toxin consists of three bacterial derived proteins, the lethal-factor (LF), the edema-factor (EF), and the protective antigen (PA). The two cytotoxic enzymes LF and EF are delivered into the cytoplasm of the intoxicated cell by the PA, which undergoes multistep structural transformations to carry out their translocation into the cell.

After the proteolytic activation by the host’s proteases, the PA chains self-assemble forming the prepore ring-like oligomer which consists of 7-8 subunits [4–6]. The prepore recruits the LF and EF and mediates the toxin’s transfer into the cell by docking on the host’s cell surface receptors, followed by endocytosis [4,7–10]. Upon endosome acidification, the prepore undergoes a cooperative structural transition forming an elongated beta-barrel hose-like structure that opens a pore in the endosome’s membrane. Through this pore, the LF and EF are extruded into the cytoplasm [11–13].

PA is composed of four domains, D1-4, each with a distinct functional role. The apical D1 domain mediates the interactions with LF and EF. The D2 domain, which is mostly sequestered inside the prepore ring-like structure, forms the membrane penetrating beta-barrel upon pore formation. The D3 domain provides most of the inter-subunit interactions that stabilize the prepore homo-oligomer. The bottom facing D4 domain mediates the interactions between the PA and the receptors on the plasma membrane of the intoxicated cell (Figure 1).

**Figure 1:**
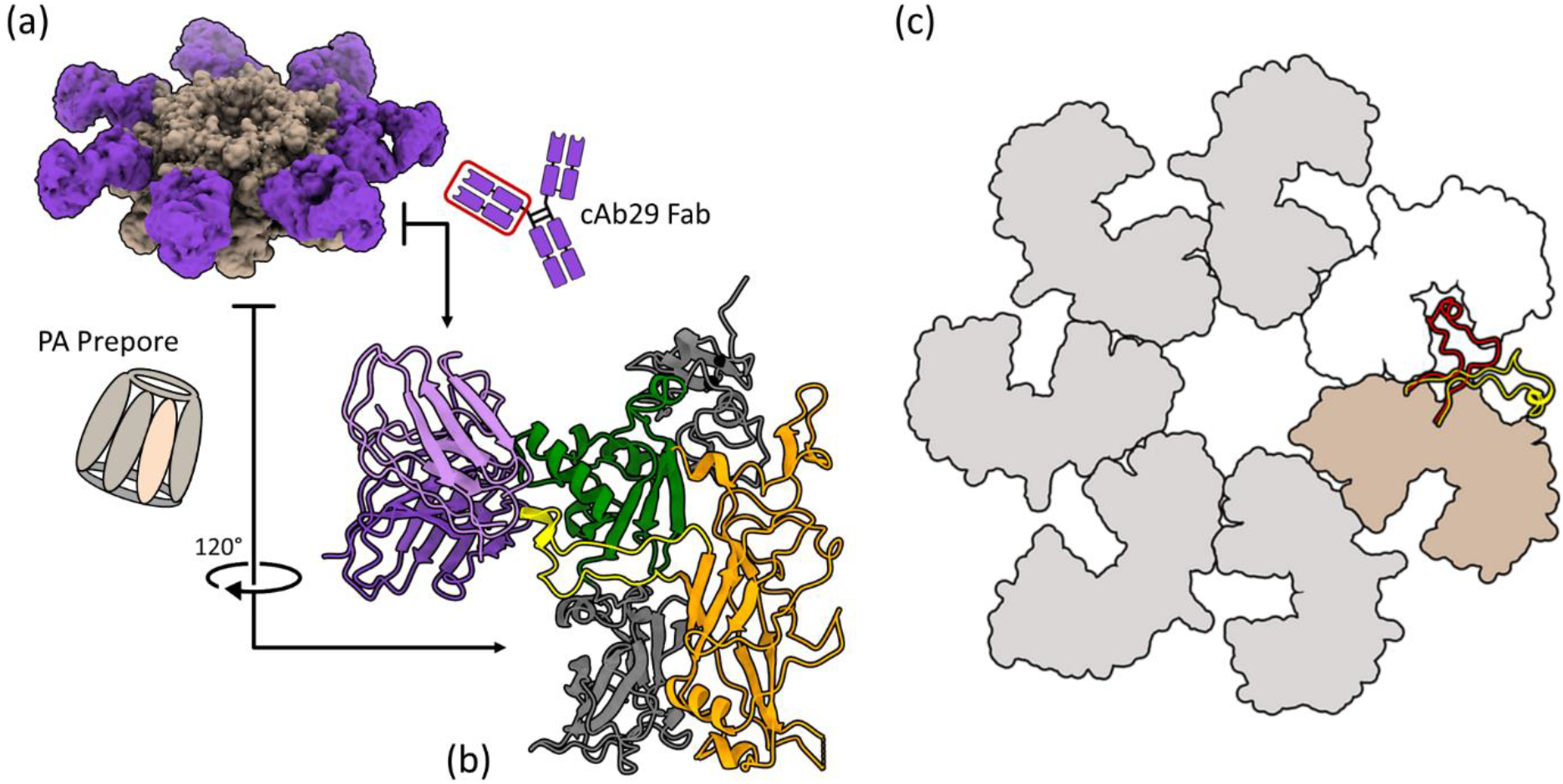
Overall structure of the PA-cAb29 complex. (a) Segmented cryo-EM density map of PA (tan) in complex with the cAb29-Fabs (violet). (b) A ribbon diagram of the PA-cAb29 complex, showing the position of the D2L2 loop (yellow), at the interface between cAb29 (light and dark violet, for the light and heavy chains respectively) and the D3 domain (green). The rest of the D2 domain is orange. (c) A slice through the top view of the prepore showing silhouettes of four neighboring subunits. Only the D2L2 loop of the tan subunit is presented, using a ribbon representation. In the cAb29 bound conformation the D2L2 loop (yellow) it is folded towards its own (tan) PA subunit, while in the cAb29-free structure (PDB ID 1TZO) the D2L2 loop (red) is swept into the inter-subunit cleft towards its neighboring subunit (white).

Due to its role in the pathogenicity of *B. Anthracis* and its multistep mechanism of action, PA is the target of monoclonal-antibodies (mAb) based therapy [14]. cAb29 is an exceptionally effective mAb anthrax-toxin neutralizer [15,16], which inactivates the toxin by preventing the prepore to pore transition [15]. To reveal how cAb29 abrogates the prepore-to-pore transition, we solved the structure of the prepore in complex with the fragments-antigen-binding (Fabs) derived from cAb29. We found that cAb29 binds the D2L2 loop that forms the membrane penetrating hydrophobic rim in the pore conformation.

Based on the structural analysis, we propose that cAb29 neutralizes the anthrax toxin by conformational selection, capturing PA’s membrane penetrating loop while it is in its solvent-exposed state and stabilizing it in a conformation that cannot form the pore.

## Results

### The overall structure of the PA-cAb29 complex

To understand how cAb29 binds PA and prevents pore formation (Figure S1 and [15]), we solved the structure of the PA-cAb29-Fab complex (PDB ID 7O85) to a resolution of ~3.3 Å, using cryo-EM. The overall shape of the prepore was similar to the heptameric prepore structures found in previous structural studies of PA (PDB ID 1TZO) [4] (Figure 1). The Fabs’ electron densities were found near the D3 domains of the prepore oligomer, arranged as seven symmetric petals with nearly coplanar alignment between the PA ring and the Fabs. 2D classification revealed a small fraction of octameric PA oligomers. The arrangements of the cAb29-Fab around both the octameric and heptameric prepores were similar, suggesting the same binding site for the eight and seven membered PA rings (Figure S2b-c).

### The interaction of cAb29 and PA

The Fab’s position near the top part of the D3 domain was unexpected. This region of the prepore does not participate in the prepore to pore transition, and its structure is almost unaffected by this transition [13]. Therefore, it was not immediately clear how the binding of cAb29 to that region could abrogate the prepore to pore transition.

Further examination of the structure revealed that in the PA-cAb29 complex, the PA’s membrane penetrating loop (D2L2 loop) is dislodged from the inter-subunit cleft formed between its own and its neighboring subunits (PDB ID 1TZO [4]). Instead, the D2L2 loop is folded away from its neighboring subunit and towards the top part of the D3 domain (Figures 1b-c, 2). As a result, all the loop’s interdomain interactions are severed, and new intradomain interactions are formed (Figure 2a). About two-thirds of the Fab’s antigen-binding surface (~660 Å^2^) interacts with the D3 domain, and the remaining surface (~338 Å^2^) interacts with the D2L2 loop (Figure S3) [17].

**Figure 2:**
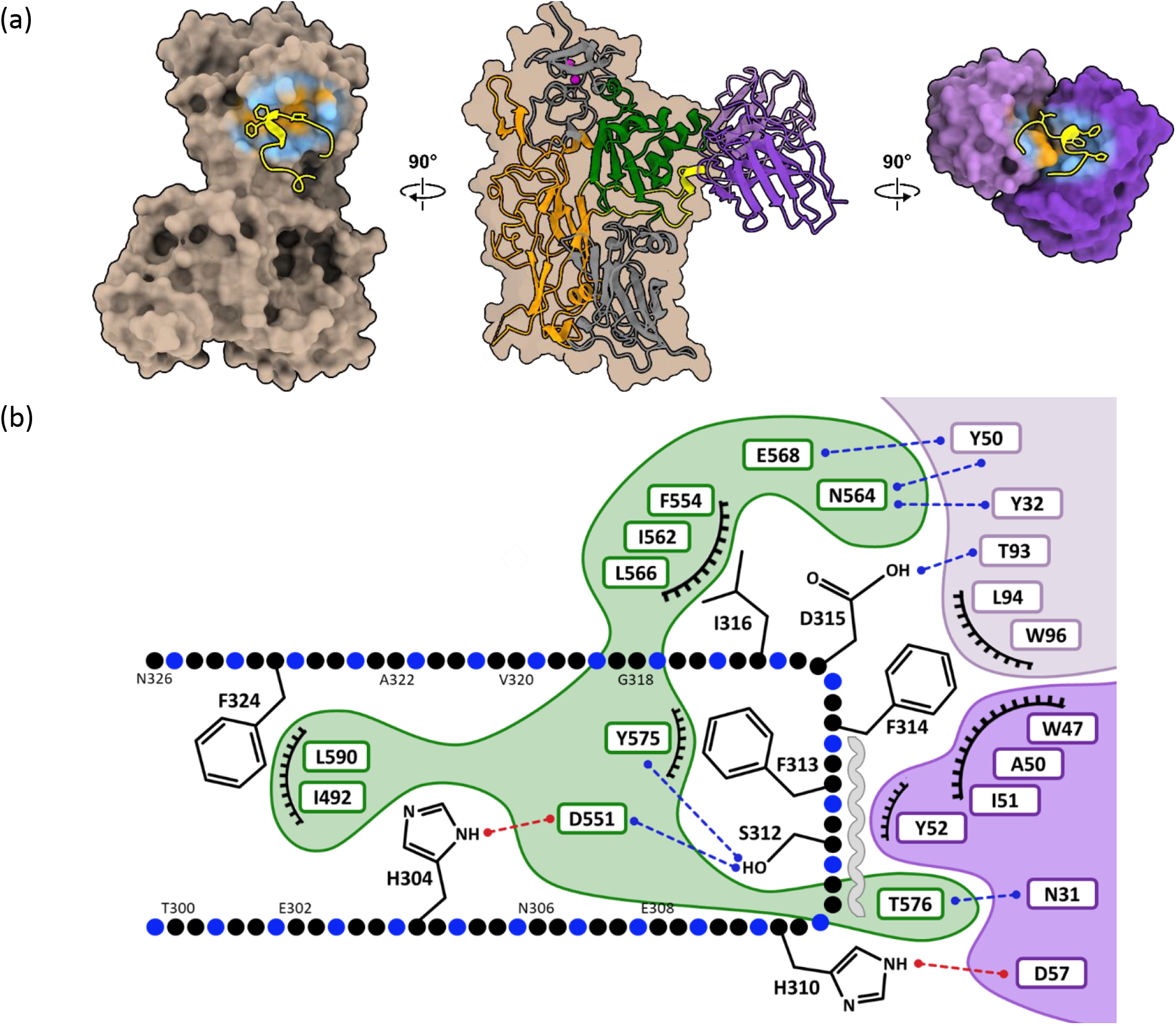
Interactions between the D2L2 loop, cAb29 and the D3 domain. (a-center) The overall domain organization of a single PA subunit (D1 and D4 – gray, D3 – green, D2 – orange, and the D2L2 loop – yellow), and its interactions with the antigen-binding segment of cAb29 (light and dark violet for the light and heavy chains, respectively). (a-left and right) Rotating PA and cAb29 by 90 in opposing directions exposes their interface. The D2L2 loop is depicted on each surface by a yellow ribbon representation. PA and cAb29 are presented in space filling surface representation. PA is tan and the light and heavy chains of cAb29 are light and dark violet, respectively. The interfaces are colored by hydrophilic-hydrophobic scale (light-blue to orange), revealing the hydrophobic pockets which harbor the loop’s hydrophobic residues and the hydrophilic rings surrounding them. (b) The interconnected surfaces of the D3 domain (green background), the light and heavy chains (light- and dark- violet backgrounds, respectively) and the D2L2 (pearl representation) presented in a 2D diagram showing the main interacting amino acids. Hydrogen bonds are blue, salt bridges are red and interacting hydrophobic regions are presented by eyelashes. The helical segment of the D2L2 loop is marked by a gray wave.

In its new Fab-bound conformation, the D2L2 loop is confined from all sides (Figure 2). The outward-facing surface of the loop is bound to the Fabs’ CDRs, and its inward-facing surface is embedded into a shallow indentation on the prepore’s side. Two hydrophobic pockets at the center of the shallow indentation in the PA’s surface harbor the highly hydrophobic membrane penetrating segment of the D2L2 loop F313-F314 in one pocket, and I316 in the other. The pockets are capped by hydrophobic thumbs (I51 and L94) on the Fab (Figure 2b). Short-range rearrangement inside the PA involving Y575 and F552, and the occlusion of F313 inside one of the hydrophobic pockets, place these aromatic residues in a three-layered staggered π - π stacking motif, further contributing to the stability of this conformation. Two rings of polar residues, one on the PA side and the other on the Fab side, surround the hydrophobic pocket and seal it (Figure 2a left and right, and Table S1).

### The conformation of the membrane penetrating loop in various structures

In most PA structures, the stems of the D2L2 loop are resolved (e.g., up to S301, and from S325) while the central part of the loop is not (Figure S4). In the few structures where the D2L2 loops are complete [4], these loops consist of a short central helical segment (A311-F314) flanked on both sides by glycine rich coils. Alignment of these structures, along with the structures where the loops are only partially resolved, reveals two main hinges. One hinge (S301) on the N-terminal side of the loop and another (G323) on its C-terminal side (Figure S4a). Comparing the orientations of the stems of the loop positioned between the main hinges can provide a rough indication of the central segment’s bearings. Thus, the rough position of the loop’s central segment, i.e., towards the D4 domain and the inter-subunit cleft, or the D3 domain and the solvent can be deduced even when this central segment of the D2L2 loop is missing. Based on this approach, we analyzed all the PDB structures of PA in the prepore and monomeric states [4,8–10,18–24] and compared the orientations of the stems of the D2L2 loops in all of these structures. In some of these structures, the stems are not resolved past the main hinges and therefore they do not provide any clues regarding the position of the loops’ central segment (Figure S4b gray). However, in other structures, the orientations of the chain in the segments flanked by the main hinges points away from the D4 domain (Figure S4 purple), while other point towards it (Figure S4 red); indicating that the D2L2 loop can adopt a wide range of conformations, some of them are solvent exposed.

## Discussion

Understanding why certain antibodies neutralize their target while others bind it without causing neutralization is crucial for developing effective neutralizing antibodies and more efficient vaccines. To provide additional insights to this central question, we set out to examine the structural mechanism of the anthrax toxin’s inactivation by a highly potent neutralizing antibody. cAb29 is an effective neutralizer of anthrax toxin, which caught our attention for its ability to block the prepore to pore transition. We assumed that cAb29’s function could be explained either by interfering with the pH-sensing of the prepore, as suggested by earlier research [15,25], or by interacting with PA’s membrane penetrating loop, thereby directly preventing its movement.

We solved the structure of the PA prepore in complex with cAb29 to determine its neutralization mechanism. We found that cAb29 inhibits the prepore to pore transition by binding to the D2L2 loop’s membrane penetrating region and clamping it to the apical part of the D3 domain (Figures 1-2). This new conformation of the D2L2 loop is stabilized by its interactions with PA’s surface and the Fab. The exact match between hydrophobic regions on PA’s surface and the hydrophobic elements of the D2L2 loop suggests that this conformation also occurs without the Fab. Therefore, it is likely that cAb29 binds the D2L2 loop while sampling exposed conformations, suggesting a conformational selection mechanism.

Several findings support a conformational selection mechanism for the binding of cAb29 to PA. Limited proteolysis experiments of the prepore lead to the cleavage of the D2L2 loop, indicating that in the prepore state the loop reversibly visits solvent exposed conformations [26]. In contrast, in the prepore’s crystal structures (PDB ID 1TZO), the D2L2 loops are swept into a cleft formed between neighboring PA subunits [4]. Similarly, in three out of seven subunits of the asymmetric PA complex (PDB ID 6UJI), the D2L2 loops reside in the inter-subunit cleft. However, the D2L2 loops in the other four subunits are unresolved [25]. In all crystal structures of monomeric PA, the D2L2 loops are absent as well [8,18–22,24,27]; this is also true for the recently published cryo-EM structure of the prepore-LF complex that otherwise has an excellent resolution [9]. Therefore, in the prepore state, the D2L2 loop samples a broad conformational spectrum ranging from solvent sequestered to solvent exposed positions.

To further establish the flexible nature of the D2L2 loop in the monomeric and prepore states, we consolidated PA’s extensive body of structural data comparing the conformations of the loops’ stems. Alignment of PA structures from the PDB (see the methods section for a complete list) showed that the loop’s stems’ orientations vary between several positions, indicating that the loop adapts a wide array of orientations. (Figure S4). Taken together, cAb29 stabilizes an exposed conformation that is sampled by the D2L2 loop without the antibody. Consequently, our results provide a mechanistic basis for using the D2L2 as a target for developing anthrax vaccinations aimed at eliciting the formation of effective neutralizing antibodies [14,28,29].

The majority of the interactions between cAb29 and the prepore are with the D3 domain, which contrasts with the dynamic nature of the D2L2 loop, which is structurally stable (Figures 2). These interactions constitute the polar rings (Figure 2b), which set the stage for forming the hydrophobic pocket that harbors the D2L2 loop. Therefore, it is likely that the PA-cAb29 interaction starts by binding cAb29 to the D3 domain, followed by stabilizing the D2L2 loop in its new conformation, when it samples conformations outside the inter-subunit pocket.

Taken together, our results provide the following structural mechanism for the inactivation of anthrax toxin by cAb29, a D2L2 loop-binding antibody (Figure 3). In the prepore state, the D2L2 loop, including its hydrophobic membrane penetrating segment, intermittently samples conformations outside the cleft between neighboring PA subunits (Figure 3 bottom left panel). When exposed, the membrane penetrating segment of the D2L2 is captured and fastened to the apical region of the D3 domain, which serves as an initial anchoring point for the cAb29-PA interactions (Figure 3 top panel). Capturing the D2L2 loop by cAb29 inhibits pore formation, thereby blocking the delivery of the cytotoxic enzymes from the endosome to the cytoplasm. As a result, the entire toxin assembly, including PA and the cytotoxic enzymes LF and EF, remains endosome locked.

**Figure 3:**
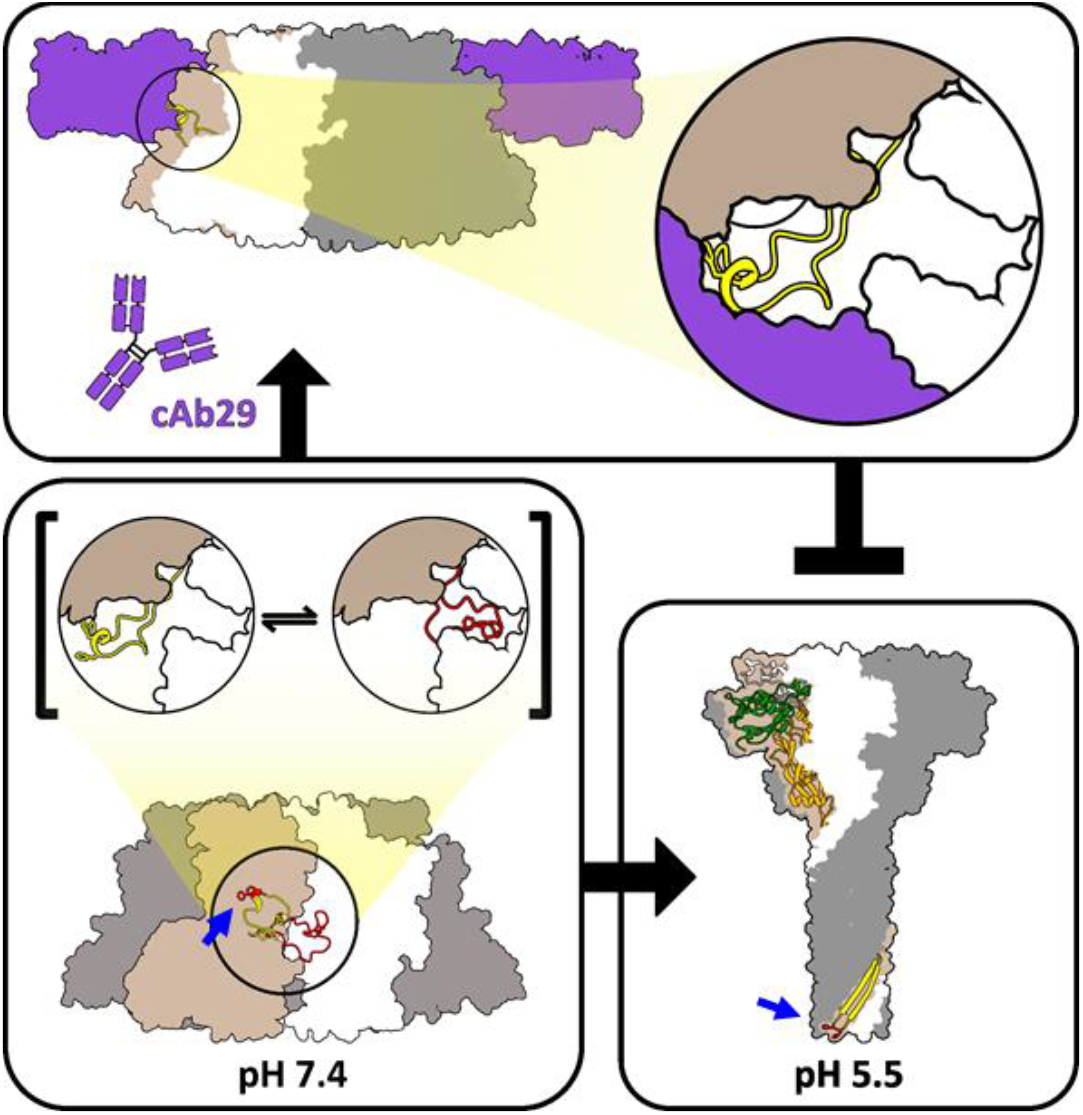
Structural mechanism for the neutralization of the anthrax toxin by cAb29. In the prepore state (bottom left panel pH 7.4) the D2L2 loop dynamically visits exposed conformations. When freely moving and upon activation by the pH drop in the endosome, the D2L2 loop can assume its position at the rim of the pore (bottom right panel pH 5.5). (c) cAb29 binds to the D3 domain and captures the D2L2 loop (top panel). When clamped to the D3 domain, the D2L2 cannot undergo its acidity induced conformational change. As a result, the PA pore is not formed and the toxin components remain endosome locked. The markedly different position of the membrane penetrating segment of the D2L2 loop in prepore (bottom left) and the pore (bottom right) are designate by blue arrows. Enlarged segments with top view orientations are marked by circles.

## Methods

### Preparation of cAb29 Fab

Chimeric anti-PA monoclonal antibody (cAb29) was produced in CHO cell line that is stably express the antibody [16], purified by affinity chromatography on HiTrap Protein A (GE Healthcare, UK) and dialyzed against PBS pH 7.4. Next, The cAb29 was concentrated to 8 mg/ml using Amicon Ultra Centrifugal Filter Unit (Merck, USA) followed by buffer exchange to 20 mM Tris-HCl pH 8.0 (Buffer A), using a Zeba Spin desalting column (Thermo Fisher Scientific, USA). Antibody digestion was performed using a Fab preparation kit (Thermo Fisher Scientific, USA) and the resulting Fab fragments were analyzed using SDS-PAGE. To validate that the purified Fab has retained its activity, ELISA was performed using PA as the coating antigen (Figure S1).

### Prepore formation and purification

PA was purified from the supernatant of *B. anthracis* strain V770-NP1-R (ATCC 14185) as described previously [30]. To create the prepore, the PA was nicked by incubation with trypsin for 30 min at room temperature and the reaction was stopped by the addition of a 5-fold excess of soybean trypsin inhibitor ([15]. The nicked PA then was desalted, loaded on a HiTrap Q-column (GE Healthcare) in a buffer A and the prepore fraction was eluted by 1 M NaCl gradient followed by analysis on SDS-PAGE (Figure S1).

### Measurement of cAb29 and cAb29-Fab affinities to the prepore

Direct ELISA was performed to evaluate the binding of cAb29 and cAb29-Fab to PA. Maxisorp 96-well microtiter plates (Nunc, Roskilde, Denmark) were coated overnight with 5 μg/ml of PA in NaHCO_3_ buffer (50 mM, pH 9.6), washed, and blocked with PBST buffer (0.05% Tween 20, 2% BSA in PBS) at room temperature for 1 h. Antibodies diluted in PBST were then added for 1 h, washed and incubated with detection antibody (AP-conjugated Donkey anti-human IgG (Jackson ImmunoResearch, USA, Cat# 709-055-149, lot 130049, used at 1:1000) and further developed with PNPP substrate (Sigma, Israel, Cat# N1891). Data was fitted using non-linear regression by Prism software version 8 (GraphPad Software Inc., USA) and presented as percent of B_max_.

### Sample preparation

PA-cAb29 complex was prepared by incubation on ice of freshly thawed PA prepores with cAb29 Fab for 10 min, at a final concertation of ~5 μM-monomers and ~6 μM, respectively, leading to a ratio of ~9 Fab molecules per 7 PA chains. After incubation, the reaction mix was concentrated by a centrifugation filtration device to a final concentration of ~1.5 mg/ml PA-cAb29 complex solution. All concentrations were assessed by UV280 absorbance and verified by SDS-PAGE gels.

Immediately before grids preparation, the PA-cAb29 complex solution was supplemented with DDM to a final concentration of 0.075 mM (to mitigate the strong preferred orientation formed in these samples). Grids were prepared by plunge freezing into liquid ethane using a home-built plunge freezing device. 3 μL of the PA-cAb29 complex were applied to the surface of each freshly glow-discharged Quantifoil R1.2/1.3 holey carbon grids (Quantifoil Micro Tools GmbH, Germany). Prior to plunge freezing, grids were blotted manually from the back for ~4 s. The frozen grids were stored in liquid nitrogen until imaging.

### Data collection

Samples were loaded under cryogenic conditions and imaged a Tecnai F30 Polara microscope (FEI, Eindhoven) operated at 300 kV, equipped with a K2 Summit direct electron detector fitted behind an energy filter (Gatan Quantum GIF) set to ±10 eV around the zero-loss peak. The calibrated pixel size at the sample plane was 1.1 Å. 50 frames dose fractionated micrographs were collected in a counting mode at a dose rate of ~8 e-/(pixel·s) and a total dose of ~80 e-/Å^2^. Low dose data collection was managed by a homemade automated SerialEM [31] data collection script [32], set to collect at random defocus values at a range of −1.0 to −2.5 μm.

### Image processing and model building

Movies were motion corrected and dose-weighted using MotionCor2 [33] and defocus values were estimated using the CTFfind4 [34]. Particle were picked followed by several iterations of 2D classifications by cryoSPARC [35–37], done for removing falsely picked carbon, ice particles, and contaminations. Further 2D classification was done for removing octameric particles (Figure S2), followed by preparation of a 3D initial model assuming C7 symmetry, this map was used for the generation of the low-resolution figure of the overall structure (figure 1a). Image processing was continued with Relion 3.0 [38]. Particle coordinates were transferred from the cryoSPARC 2.15 database to Relion’s *.star format following Daniel Asarnow’s script (csparc2star.py, https://github.com/asarnow/pyem/wiki/Export-from-cryoSPARC-v2) [35–37]. Using Relion, we performed an additional round of 2D classification followed by the selection of 2D classes with class averages showing the clearest feature. These particles were then refined, post-processed and polished following Relion’s pipeline. The main stages of the image processing are documented in Figure S2 and the details of data collection and image processing are tabulated in table S2.

The model (PDB ID 7O85) was built by several iteration manual model building in coot followed global real space refinement using Phenix [39], starting from the prepore crystal structure PDB ID 1TZO [4], and a Fab model PDB ID 4ZSO.

### Multiple structure alignment

The coordinates of the following structures were fetched using Chimera [40]: 4H2A 1T6B 6ZXJ 6ZXK 6ZXL 3Q8A 3Q8F 3MHZ 3Q8B 1ACC 4EE2 4NAM 3Q8E 3Q8C 3TEZ 6WJJ 6VRA 5FR3 3TEX 3TEW 3TEY 1TZO 1TZN 3KWV 3HVD. The structures were split to monomers, keeping only one chain for each symmetric structure and deleting non-PA chains (e.g., the chain of UMG2). All amino acids up to amino acids 258 and from amino acid 488 up to the end of the sequence were deleted, leaving only the structures of the D2 domain (i.e., amino acids 258-487). The structure of the all the D2 domains were aligned to the structure of PA determined in this study using the match align command of chimera. The main hinges were located by manual inspection looking for the point where the different chains diverged. After locating the main hinges, the electron density cloud (when available) corresponding to the different structures were manually inspected in order ensure that the PDB structure and the electron density maps are in good agreement at the region we were analyzing. Jalview 2.11.1.4 [41] was used to automatically generate a multiple sequences alignment of all the D2L2 loops marking the absent parts of the structures as gaps. This alignment was used for determining the structures that have at least two amino acids resolved past the main hinges of the D2L2 loops.

## Supporting information

Supporting experimental results and additional figures and tables

## Acknowledgments

The research was supported by the Agreement on Industrial, Scientific and Technological Research and Development Cooperation between the Israeli Ministry of Science, Technology and Space (MOST) and the Ministero Italiano degli Affari Esteri e della Cooperazione Internazionale (MAECI) (Project: SB2D-CNI-BGU); and by the ISRAEL SCIENCE FOUNDATION (grant No. 364/20). The authors wish to thank Mr. Yehuda Barouch for IT support, Prof. Anat Ben-Zvi for constructive criticism of the manuscript, and Dr. Stephen H. Leppla for useful advice and discussions.

