## Supporting experimental results and additional figures and tables for "Neutralization of the anthrax toxin by antibody-mediated stapling of its membrane penetrating loop"

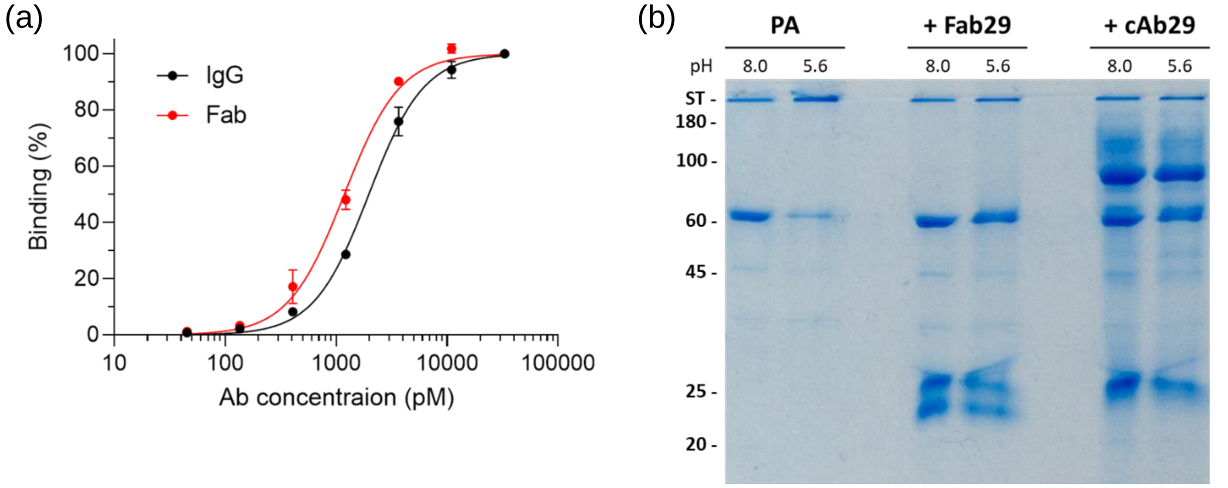


**Figure S1: cAb29 and its Fab bind the prepore and block the prepare to pore transition. (a) Binding profiles of cAb29 antibody and cAb29-Fab tested by ELISA against PA. Values are averages of duplicates ± SEM, fitted by non-linear regression and expressed as percent of Bmax of each curve. The dissociation constants of the Fab and the antibody are 1.97±0.06 nM and 1.18±0.07 nM respectively. (b) Transition to the pore state was monitored by the formation of an SDS-resident PA complex which did not migrate into the separating gel. The ability of PA to undergo the transition was tested by incubating PA, w/wo cAb29 or cAB29-Fab at a ratio of 8:7 Fab:PA-subunits, for 15 min, followed by an additional 15 min incubation at pH 5.6 (or pH 8.0, as a control), both incubations were done at room temperature. After incubation in low pH, SDS-PAGE samples were prepared by adding Laemmli buffer to the PA samples to a final concertation of 1x followed by incubation 10 minutes in room temperature prior to gel loading. Samples were resolved on 10% acrylamide gel. Samples' pH before addition of SDS is noted above the lanes. Only when the Fab and IgG are absent, treatment with low pH resulted in a decrease of the SDS sensitive phase manifested as a band expected of monomeric $PA_{63}$ ($2^{nd}$ lane, 63 kDa band).**

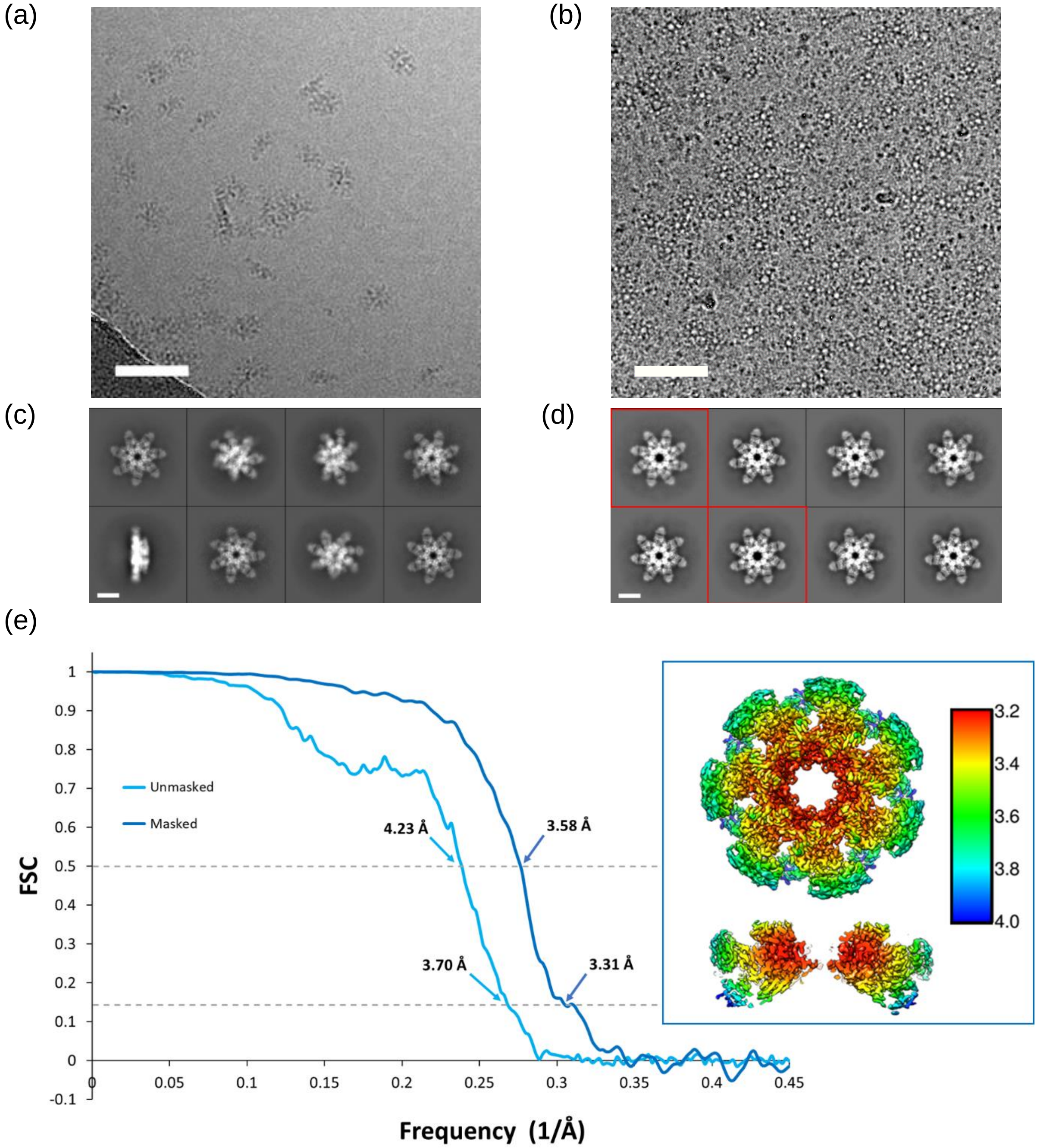


**Figure S2: Summary of image processing of the cAb29-PA complex datasets. (a-b) Representative micrographs (cropped to ~0.25 field of view) of cAb29-PA complexes w/wo addition of 0.075 mM DDM (respectively). (c-d) Selected "good" 2D class averages form the dataset w/wo DDM (respectively). Class averages of top views of the octameric complex are highlighted by a red frame. Addition of DDM to the sample resulted in a broad orientation distribution of the PA-Fab complexes, however reduced the number of particles per field of view. (e) FSC plot of the resulting 3.5 Å map and the corresponding local resolution maps show a top over view of the structure and a slice through its center (inset). Scale bar in the micrographs is 50 nm and on the 2D class averages is 10 nm.**

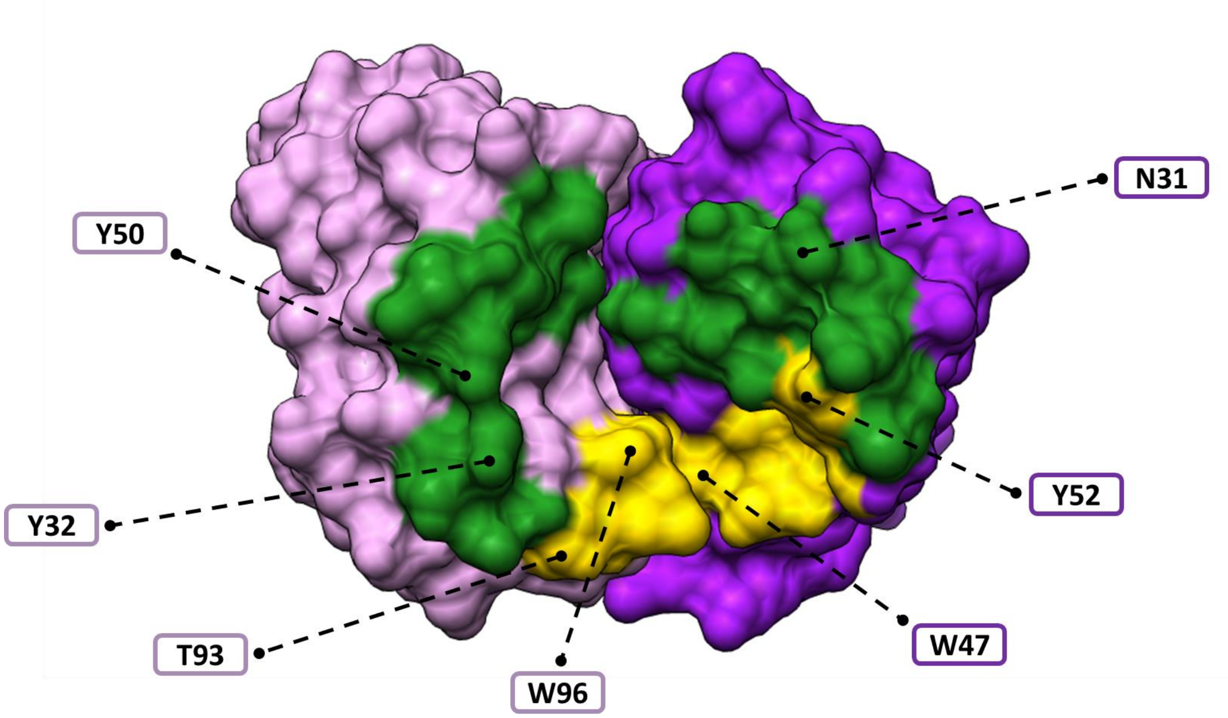


**Figure S3: The footprint of the D2L2 loop and the D3 domain on the antigen binding surface of cAb29. The light and heavy chains of cAb29 are colored light and dark purple, respectively. The surface of cAb29 that interacts with the D2L2 loop is colored yellow and the surface that interacts with the D3 domain is colored green. The position of some of the amino acids which interact with PA are designated (see figure 2b for a detailed interaction plot). The cAb29-Fab antigen binding surface is exposed by removing the PA and looking outwards from the prepore ring towards the Fab.**

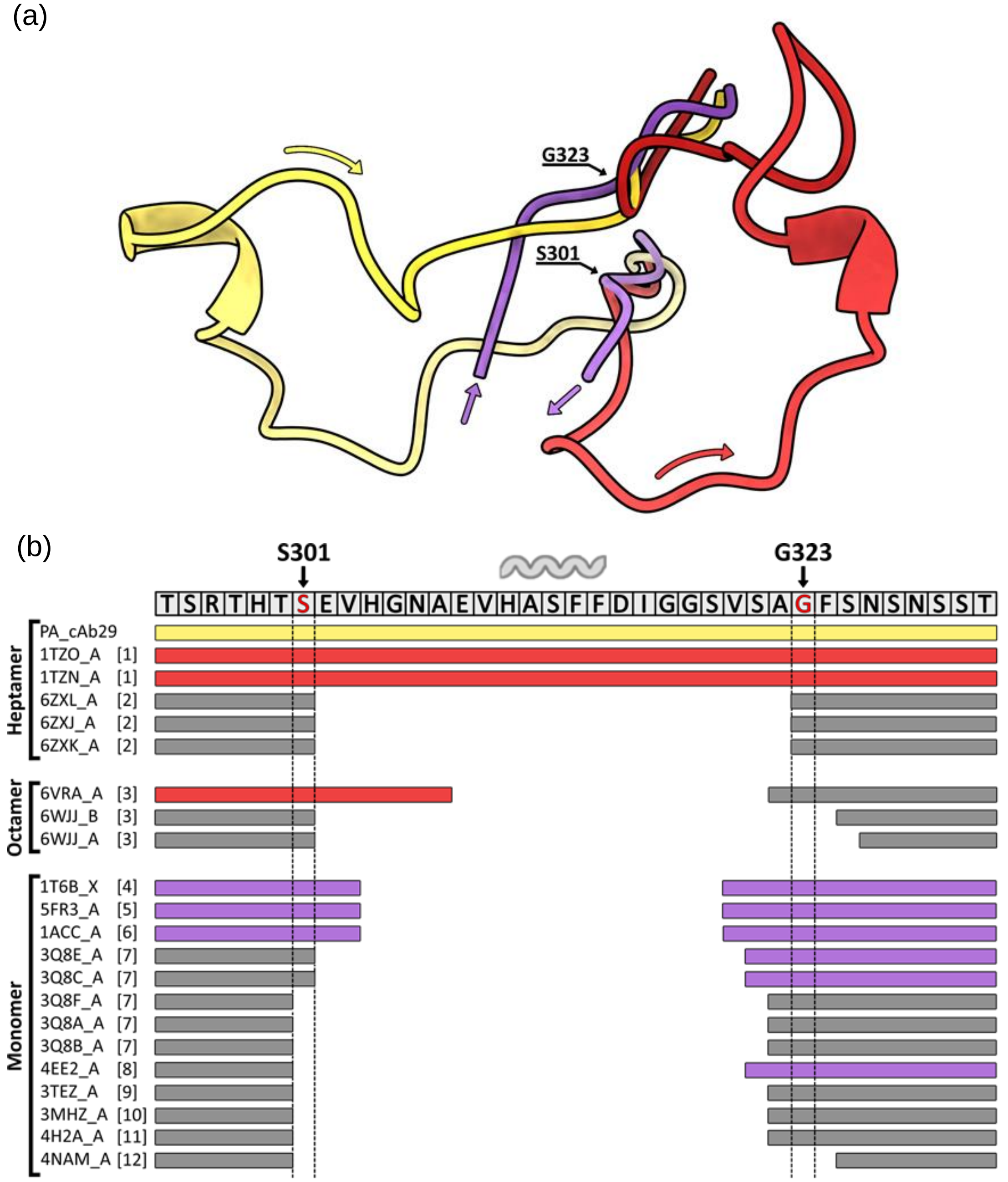


**Figure S4: The D2L2 loop has a wide conformational spectrum. (a) Alignment of 22 structures of PA reveals the main hinges flanking the central part of D2L2 on its N- (S301) and C- (G323) terminal sides. Three representative structures are presented for clarity: the structure of the PA-Fab complex determined in this study—yellow, the crystal structure of the prepore (PDB ID 1TZO)—red, and the crystal structure of monomeric PA bound to the host cell receptor CMG2 (1T6B)—purple. Color shades in each structure designate the position along sequence from bright to dark. The rough direction of the D2L2 loop can be deduced form the direction of the loop's stems past the hinges. (b) The structurally resolved regions of the D2L2 loop in different PA structures depicted as colored bars under the loop's sequence, areas missing in the structure are empty. The bars are colored according to the similarity of each structure to the structures depicted (a). In instances where the resolved stems segment are too short (less than two amino acids past the hinge) the bars are colored gray.**

| **Table S1: List of amino acids forming the hydrophilic seal around the PA-Fab hydrophobic interface** | |
|---|---|
| **PA side** | **Fab side** |
| T548, **D551**, N556, **N564**, **E568**, T572, N573, **T576**, K594, H597 | Light chain: K92, **T93**<br>Heavy chain: **N31**, N33, R54, T55, **D57**, S59, D99 |
| * Amino acids depicted in figure 2b are in bold font | |

| **Table S2: Data collection and refinement parameters of cryo-EM structures solved in this study** | |
|---|---|
| PDB ID/EMDB Codes | 7O85/EMD-12761 |
| FSC(1/f)=0.143 Masked/Unmasked (Å) | 3.5/3.9 |
| FSC(1/f)=0.5 Masked/Unmasked (Å) | 3.8/4.4 |
| # of micrographs | 6355 |
| # of particles | 211 K |
| Total dose (e-/Å2) | 80 |
| Pixel size (Å) | 1.1 |
| # of frames | 50 |
